# Antibacterial mechanism of inosine against *Alicyclobacillus acidoterrestris*

**DOI:** 10.1101/2023.07.05.547889

**Authors:** Xiaoxue Liu, Youzhi Wu, Junjian Ran, Lingxia Jiao, Linjun Sun, Fuzhou Ye

**Affiliations:** School of Food Science, Henan Institute of Science and Technology, Xinxiang, Henan, 453003, China; School of Food and Drugs, Shanghai Zhongqiao Vocational and Technical University, 201514, Shanghai

**Keywords:** *Alicyclobacillus acidoterrestris*, Inosine, Antibacterial, Protein interaction

## Abstract

Inosine could potentially become a novel antibacterial agent against *A. acidoterrestris* as low-dose of inosine can prevent *A. acidoterrestris* contamination. However, until now the antibacterial mechanism of inosine targeting *A. acidoterrestris* is still unknown. In this study, to unravel the mechanism of inosine against *A. acidoterrestris* puzzle, the effects of inosine on bacterial surface hydrophobicity, intracellular protein content, cell membrane damage extent and permeability of the *A. acidoterrestris* were investigated. The results showed that inosine can effectively inhibit the growth and reproduction of *A. acidoterrestris* by destroying the integrity of cell membrane and increasing its permeability, causing the leakage of intracellular nutrients. Furthermore, the interaction networks of inosine target proteins were analyzed. The interaction networks further revealed that damage of bacterial cell membrane might be relevant to inosine’s effect on bacterial DNA replication and cell energy metabolism through regulating nucleotide synthesis and metabolism and the activity of translation initiation factors. Finally, the antibacterial mechanism of inosine against *A. acidoterrestris* was proposed.

## Introduction

As an acidophilic and heat-tolerant spoilage organism, *A. acidoterrestris* can produce spores, causing the spoiling of acidic fruit and vegetable products ^[1, 2]^. According to international export trade requirements, the content of *A. acidoterrestris* should not be higher than one per 10 mL concentrated juice ^[3]^. This requirement is a serious challenge to the sterilization process of juice industry. The development of a novel technique for controlling *A. acidoterrestris* contamination and improving the quality and nutrient preservation of fruit and vegetable products is crucial. At present, high-temperature sterilization can control *A. acidoterrestris* contamination well in practical production. However, there are negative effects on the nutrition, flavor and functional characteristics of fruit and vegetable products by high-temperature sterilization. The application of novel and high-efficiency bacteriostatic agents could decrease the sterilization temperature, duration, and energy consumption. Meanwhile, it increases the efficiency of equipment sterilization process and preserves the quality of fruit and vegetable products. To this point, development of high-efficiency bacteriostatic agents has a wide application in the production of high-quality fruit and vegetable.

Previous studies found that inosine is highly effective in inhibiting both *A. acidoterrestri* spore germination and bacteria growth at a concentration of 5 mM. It triggered us to think whether inosine could become a novel bacteriostatic agent against *A. acidoterrestri*. As one component of human body, inosine is environmentally friendly, colorless, and odorless, which make inosine a good candidate for controlling *A. acidoterrestris* contamination in fruit juice production. Given the crucial role of inosine in food industry, the inhibition mechanism of inosine against *A. acidoterrestris* need to be further investigated in purpose of expanding its application in food industry more safely and widely. Finally, the investigation of the antibacterial mechanism is also conducive to the development of a novel and efficient technique against *A. acidoterrestris* contamination.

This study aims to elucidate the inhibition mechanism of inosine on *A. acidoterrestris*. The parameters such as surface hydrophobicity, intracellular protein content, cell membrane damage extent and permeability *A. acidoterrestris* were measured after different concentration of inosine treatment on *A. acidoterrestris*. In addition, the proteins interaction networks of inosine target were analyzed. Taken together, this work provides further experimental and theoretical basis on developing novel controlling technique on *A. acidoterrestris* and supplying a way of antibacterial application with inosine.

## Materials and methods

### Bacterial strain and its cultivation

*A. acidoterrestris* DSM 3922^T^ was purchased from German Collection of Microorganisms and Cell Cultures GmbH (DSMZ). For sporulation of *A. acidoterrestris*, the bacterial strain *A. acidoterrestris* DSM 3922^T^ was inoculated in *Alicyclobacillus* spp. medium (AAM) prepared according to a previous study ^[4]^.

### Bacterial surface hydrophobicity determination

Cell adsorption rate was used to determine the bacterial surface hydrophobicity. Bacterial surface hydrophobicity was measured as described before ^[5]^. Briefly, 2 mL of *A. acidoterrestris* cell culture at the mid-log phase was taken. Then 0.5 mL of either 25 mM or 50 mM inosine solutions was added to make the final concentration of inosine 5 mM and 10 mM, respectively. AAM liquid medium without inosine was used as a control. After mixing, the bacterial samples were each incubated at 45°C and stirred at 250 r/min for different time duration (0, 30, 60, 90, and 120 mins separately). Another 1.5 mL of xylene was then added to each individual sample and vortexed for 30 s. The mixture was left at room temperature for another 30 mins and aspirated the lower liquid after layering. Finally, the absorbance of OD_600_ was measured by a visible spectrophotometer (SP-756PC, Shanghai spectrum instrument Co Ltd). The formula for calculating the cell adsorption rate is: Cell adsorption rate = (OD_600Control_-OD_600sample_)/ OD_600Control_×100%.

### Bacterial intracellular protein content measurement

Intracellular protein content of *A. acidoterrestris* was determined as described ^[6, 7]^. Based on previous results, the treatment duration of *A. acidoterrestris* with inosine was determined to be 6 h. As described above, *A. acidoterrestris* cultures were stirred at 250 r/min at 45°C and treated with 5 mM and 10 mM of inosine, respectively. Meanwhile, the control group was made in AAM liquid medium without inosine was examined. The samples were centrifuged to collect bacteria at 4°C (6000 ×g, 5 min). The harvested cells were suspended to about 10^6^ CFU/mL by 0.01 M PBS and then an equal amount of suspension was taken for cell lysis using ultrasonic treatment. After centrifugation, the supernatant was taken to quantify the protein content by Coomassie brilliant blue G-250 staining.

### Determination of cell membrane damage

The method to treat the *A. acidoterrestris* cultures was the same as above. According to previous work (Kang, 2021), propidium iodide (PI) was mixed with 10^6^ CFU/mL of *A. acidoterrestris* cells at a final concentration of 1 μg/mL. The mixture was then protected from light at 4°C for 30 min. After that, the mixture was washed with PBS for 3 times. Finally, 5 μL of PI staining bacterial samples was used to detect the fluorescent dye among them by an inverted fluorescent microscope (Zeiss Axio Vert A1, Germany). Additionally, another 2 mL of staining bacterial samples was analyzed with a flow cytometry to check cell viability (Beckman CytoFLEX FCM, USA, Channel FL2-A: PE-A).

### Interaction network establishment of inosine target proteins

The STITCH (http:////stitch.embl.de/) database was used to predict the interaction between inosine and gene or proteins of *A. acidoterrestris*. STITCH can predict the target of action of inosine and scored all those target relationships. A higher score indicates a higher degree of confidence ^[8]^. The inosine target gene or protein information was input in string protein interplay network database (http://www.string-db.org/) and protein interaction information for inosine target proteins was acquired. Then, the protein-protein interaction information was imported into Cytoscape to create an interaction network diagram of inosine-targeted proteins. In the diagram, nodes represent genes, molecules, or proteins and the lines between the nodes indicate the interactions between the molecules. Advance Network Merge in Cytoscape compares the interaction networks of each target protein and removes duplicate edges and isolates etc., which could obtain the largest protein interaction network of inosine target. In addition, MCODE in Cytoscape could detect densely connected regions in the largest protein network and score the connectivity of proteins. The modules with node numbers and scores both greater than 3 were selected for BinGo annotation which analyzed the biological processes participated by functional modules. The participating probability of the genes in the functional module of the biological process could be shown by p values ^[9, 10]^.

### Statistical analysis

For statistical analysis purpose, all the relevant experiments in this work are done at least 3 times repeat to meet requirement. Data were expressed as the mean ± standard deviation (SD), and Duncan’s ANOVA was analyzed using SPSS 23.0 statistical software (SPSS, Inc., Chicago, IL, USA). Fluorescence intensity data from flow cytometry detection was analyzed using FlowJo 10.0 program (Univ, Inc, Shanghai, China).

## Results

### Effects of inosine on surface hydrophobicity of *A. acidoterrestris*

The hydrophobicity of bacterial cell surface is often considered as an important regulator of cell membrane activity ^[11]^. The damage to the cell membrane can cause increased hydrophobicity of the bacterial surface. The greater the hydrophobicity, the more severe the membrane damage is ^[12]^. The hydrophobicity of bacterial surface can be judged by its xylene adsorption performance which is positively correlated with the surface hydrophobicity ^[13]^. When cell membrane is damaged, the adsorption effect of xylene is enhanced. Therefore, the change of the adsorption capacity to xylene can indicates the change of surface hydrophobicity which reflects the damage of the cell membrane ^[14]^. In our work, after inosine treatment, the adsorption capacity of *A. acidoterrestris* against xylene was significantly enhanced. It is positively correlated with the increase of inosine treatment concentration and duration (Fig. 1). When samples were treated with 5 mM inosine for 30, 60, 90, and 120 mins different time duration, the adsorption rate of *A. acidoterrestris* to xylene increased from 36.82% (in the control group without inosine treatment) to 39.33%, 43.18%, 44.91%, and 46.81% in corresponding to the four different time duration treatment. However, when the concentration of inosine increased to 10 mM to treat *A. acidoterrestris*, the adsorption rate of those 4 experimental groups increased from 36.82% (control group) to 43%, 49%, 52%, and 55%, correspondingly (Fig. 1). Taken together, these indicate high concentration and longer-term treatment of *A. acidoterrestris* with inosine, the surface hydrophobicity of cell membrane increased. These data suggest that inosine could cause damage to the cell membrane of *A. acidoterrestris*, thereby increasing the surface hydrophobicity of *A. acidoterrestris*.

**Fig. 1.**
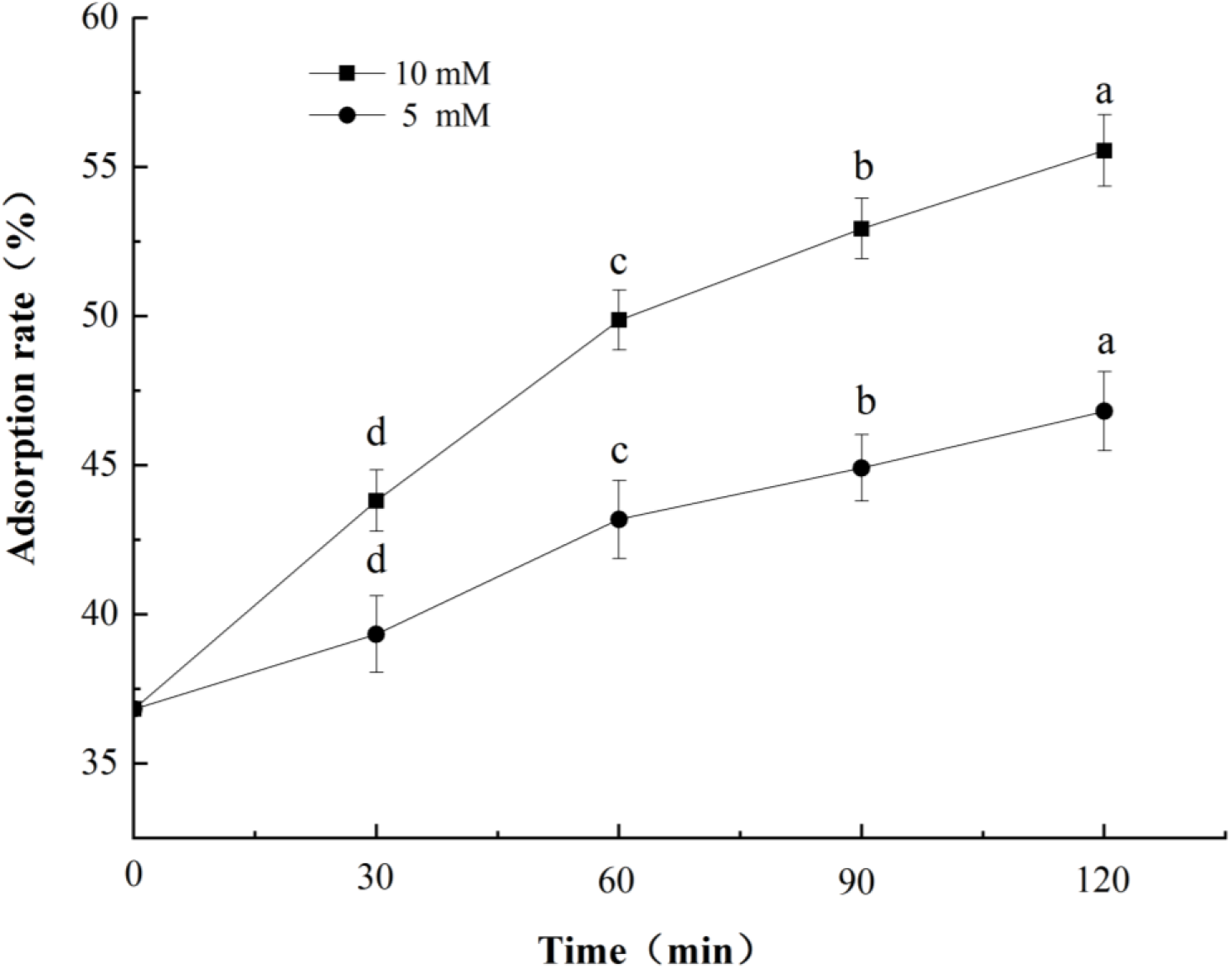
Effects of inosine treatment with different concentration and duration on *A. acidoterrestri*s surface hydrophobicity.

### Effects of inosine on intracellular proteins content of *A. acidoterrestris*

The integrity of cell membrane ensures the normal growth of bacteria. When the cell membrane is damaged, it will cause the leakage of nutrients such as intracellular proteins and result in a decrease in the intracellular protein content. Therefore, the change in the intracellular protein content can reflect the integrity of the cell membrane ^[15]^. After inosine treatment, it was found that the intracellular protein concentration of *A. acidoterrestris* decreased significantly (P<0.05, Fig. 2). When treated with 5 mM and 10 mM concentrations of inosine, the corresponding intracellular protein concentrations of *A. acidoterrestris* were 81.06 μg/mL and 68.78 μg/mL, respectively. Compared with the control group, the intracellular protein concentrations of the two experimental groups (5 mM and 10 mM inosine treatment) decreased by 46.8% and 54.9%, respectively. Higher concentration of inosine treatment caused more decrease of the intracellular protein content. This result indicates that inosine treatment caused damage to the cell membrane of *A. acidoterrestris* and resulted in its intracellular nutrition leakage.

**Fig. 2.**
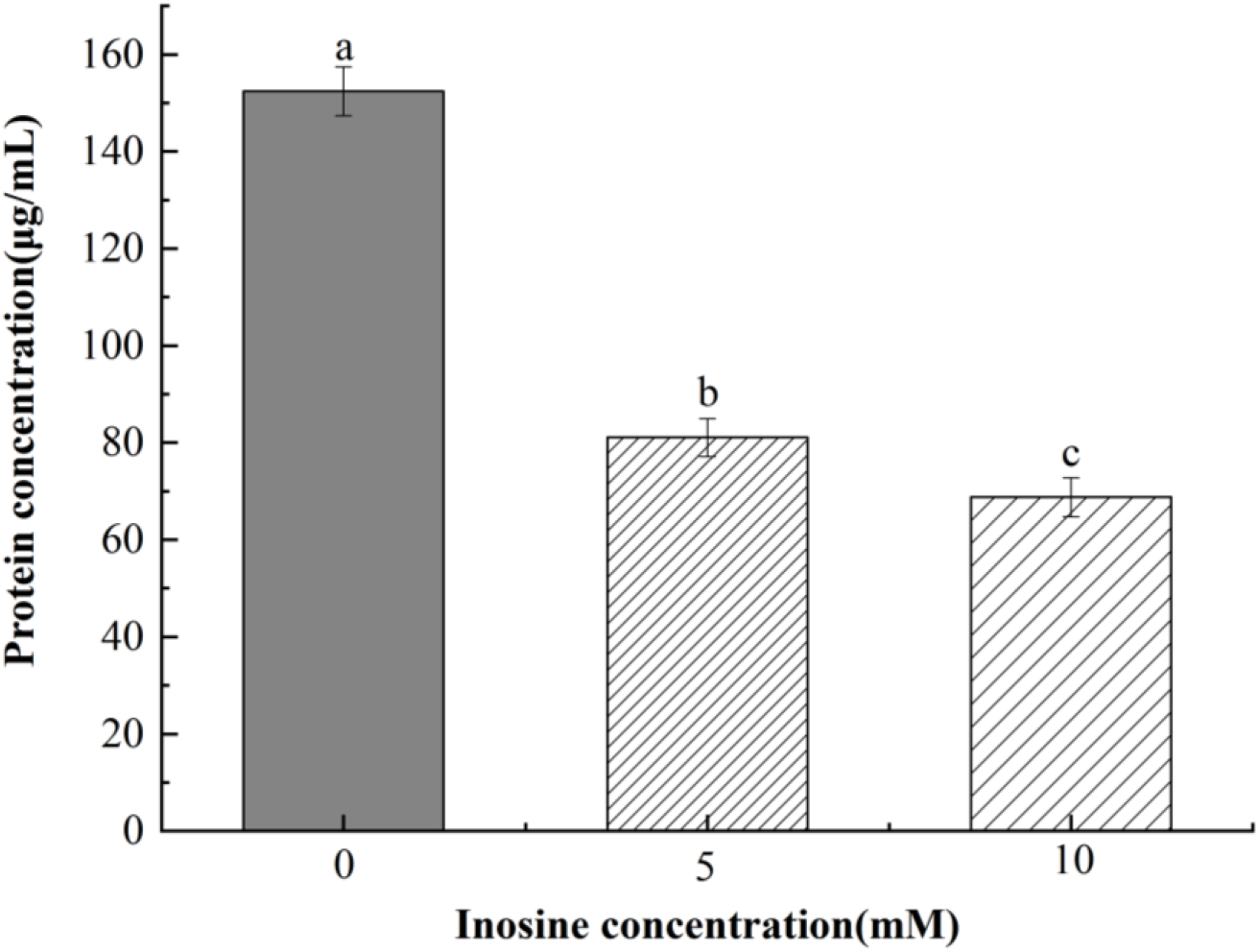
Effects of different concentration of inosine treatment on the intracellular protein content of *A*. *acidoterrestris*.

### Effects of inosine on the integrity and permeability of *A. acidoterrestris* cell membrane

Bacterial membrane damage can be detected by propidium iodide (PI) fluorescent dye labeling. Propidium iodide (PI) is a fluorescent dye commonly used for nucleic acids that do not penetrate the membranes of normal cells. However, the membranes of apoptosis cells and dead cells are destroyed so that PI can directly enter into the cells and stains the cells in red ^[6, 16]^. Therefore, the amount of red fluorescence can indirectly reflects the number of dead cells ^[17]^. After PI staining of inosine-treated *A. acidoterrestris*, fluorescence microscopy and flow cytometry were applied to detect the fluorescent intensity and population of *A. acidoterrestris*. As shown in Fig. 3A, no fluorescence was observed in the control group, while a large amount of red fluorescence appeared in the experimental group with inosine treatment (5 mM in Fig. 3B and 10 mM in Fig. 3C). Compared Fig.3 and Fig. 3B, the number of fluorescent bacteria that stained was increased with the increase of inosine concentration which indicates that inosine destroyed the integrity of the cell membrane and make it more permeable. The flow cytometry analysis shows that the main peak shifted to the right (Fig. 3D, 3E), which indicates that the intensity of fluorescence staining of *A. acidoterrestris* increases after inosine treatment which further verified the results of the fluorescence microscope detection (Fig. 3B, 3C). As shown in Fig. 3D and 3E, when *A. acidoterrestris* was treated with 5 mM and 10 mM inosine, the main peaks shifted to the right compared with the control group and the effect of the main peak shifting to the right is more significant with 10 mM inosine concentration treated. These results suggested that the integrity of the cell membrane of *A. acidoterrestris* is seriously damaged by inosine treatment, which also caused greater permeability.

**Fig. 3.**
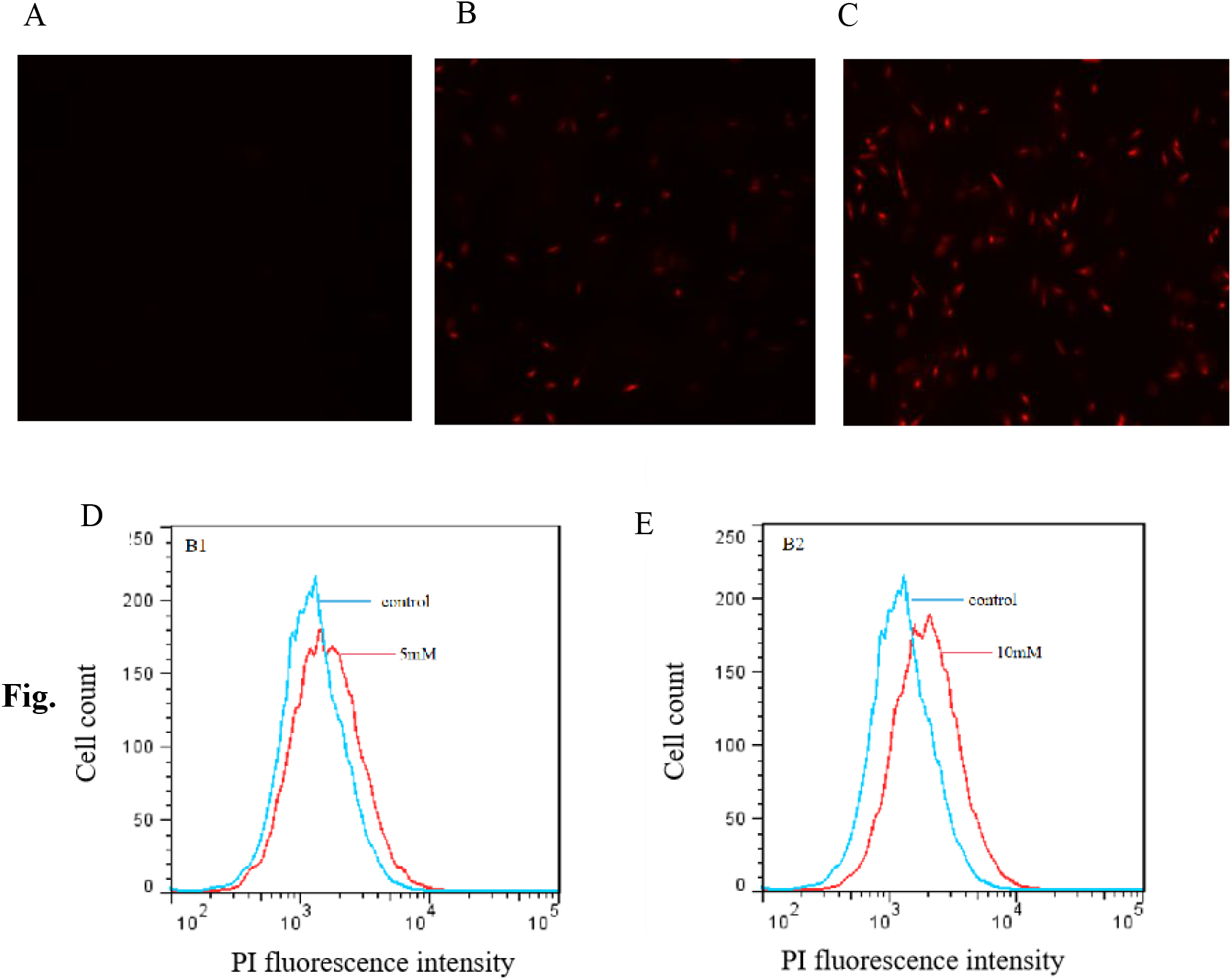
A), B), C) Micrographs of PI stained *A. acidoterrestris* cells under fluorescence microscope with different concentrations of inosine treatment. A) Control, no inosine treatment; B) 5 mM inosine treatment; C) 10 mM inosine treatment. D) and E) Flow cytometry analysis of PI stained *A. acidoterrestris* cells, after inosine treatment, D) 5 mM inosine treatment; E) 10 mM inosine treatment.

### Interaction network modules analyses of inosine target proteins

The STITCH (http://stitch.embl.de/) database can be used to predict the interactions between chemicals and genes or proteins. This study is based on a confidence level of 0.7, seven high confident target proteins that interact with inosine were searched in the STITCH database (Table 1).

**Table 1.** The information of inosine target protein.

| Targeted gene | Uniprot ID | Score | Targeted gene | Uniprot ID | Score |
| --- | --- | --- | --- | --- | --- |
| deoD | C8WUM0 | 0.982 | Aaci_0435 | C8WS60 | 0.808 |
| Aaci_0199 | C8WQT4 | 0.916 | mtaD | C8WX35 | 0.739 |
| Aaci_1647 | C8WX37 | 0.915 | Aaci_0033 | C8WPZ3 | 0.712 |
| surE | C8WX33 | 0.900 |  |  |  |

The protein information of these 7 target proteins was entered into the string database to obtain their partner proteins. In addition, the protein interaction networks of the inosine target were established using Cytoscape software, which contained a total of 54 nodes and 192 edges (Fig. 4). Using MCODE software to calculate and analyze, 5 protein interaction modules with node numbers and score values both greater than 3 were plotted (Fig. 5). By means of BinGo annotation, it was found that module 1, module 2, module 4, and module 5 were related to DNA cleavage, nucleotide metabolic process, purine nucleoside biosynthesis and translation initiation factors, respectively, while module 3 could not obtain GO annotation results (Table 2).

**Table 2.**
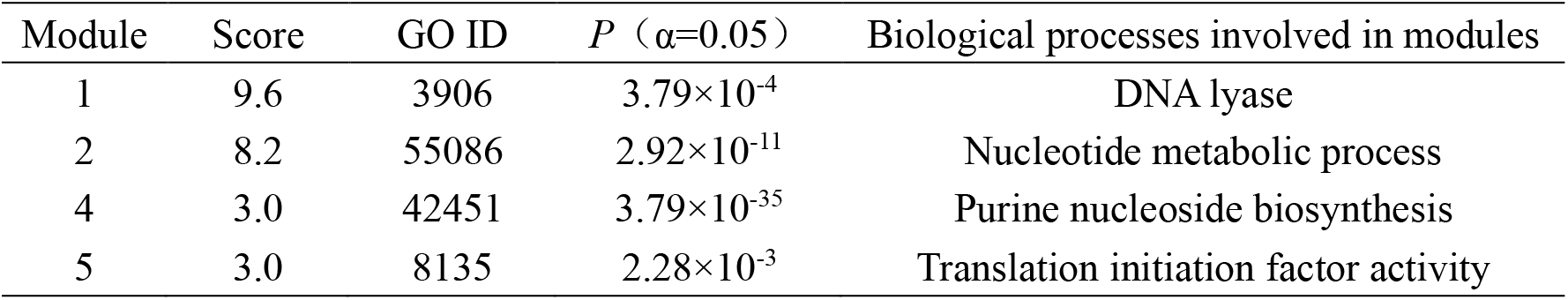
Main biological processes related to the functional modules.

| Module | Score | GO ID | $P$ ($\alpha=0.05$) | Biological processes involved in modules |
| --- | --- | --- | --- | --- |
| 1 | 9.6 | 3906 | $3.79 \times 10^{-4}$ | DNA lyase |
| 2 | 8.2 | 55086 | $2.92 \times 10^{-11}$ | Nucleotide metabolic process |
| 4 | 3.0 | 42451 | $3.79 \times 10^{-35}$ | Purine nucleoside biosynthesis |
| 5 | 3.0 | 8135 | $2.28 \times 10^{-3}$ | Translation initiation factor activity |

**Fig. 4.**
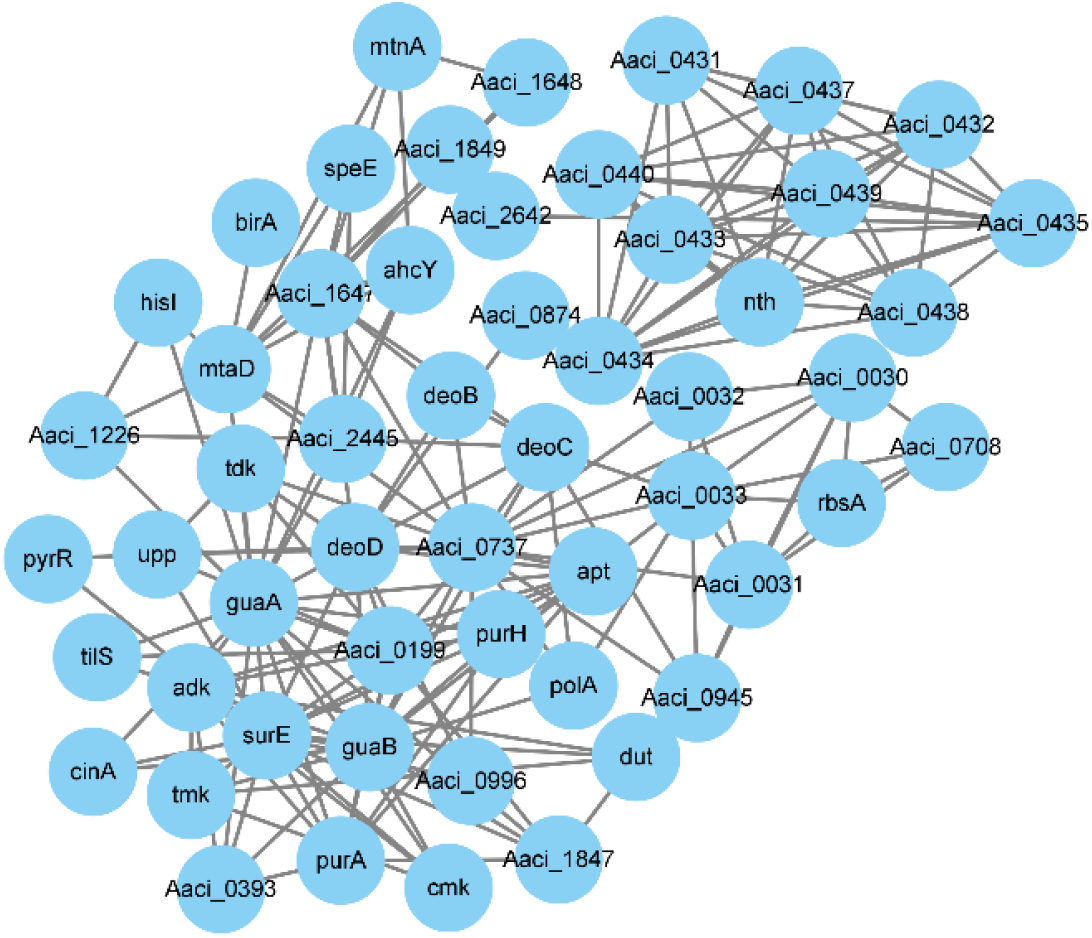
The protein interaction network of inosine action.

**Fig. 5.**
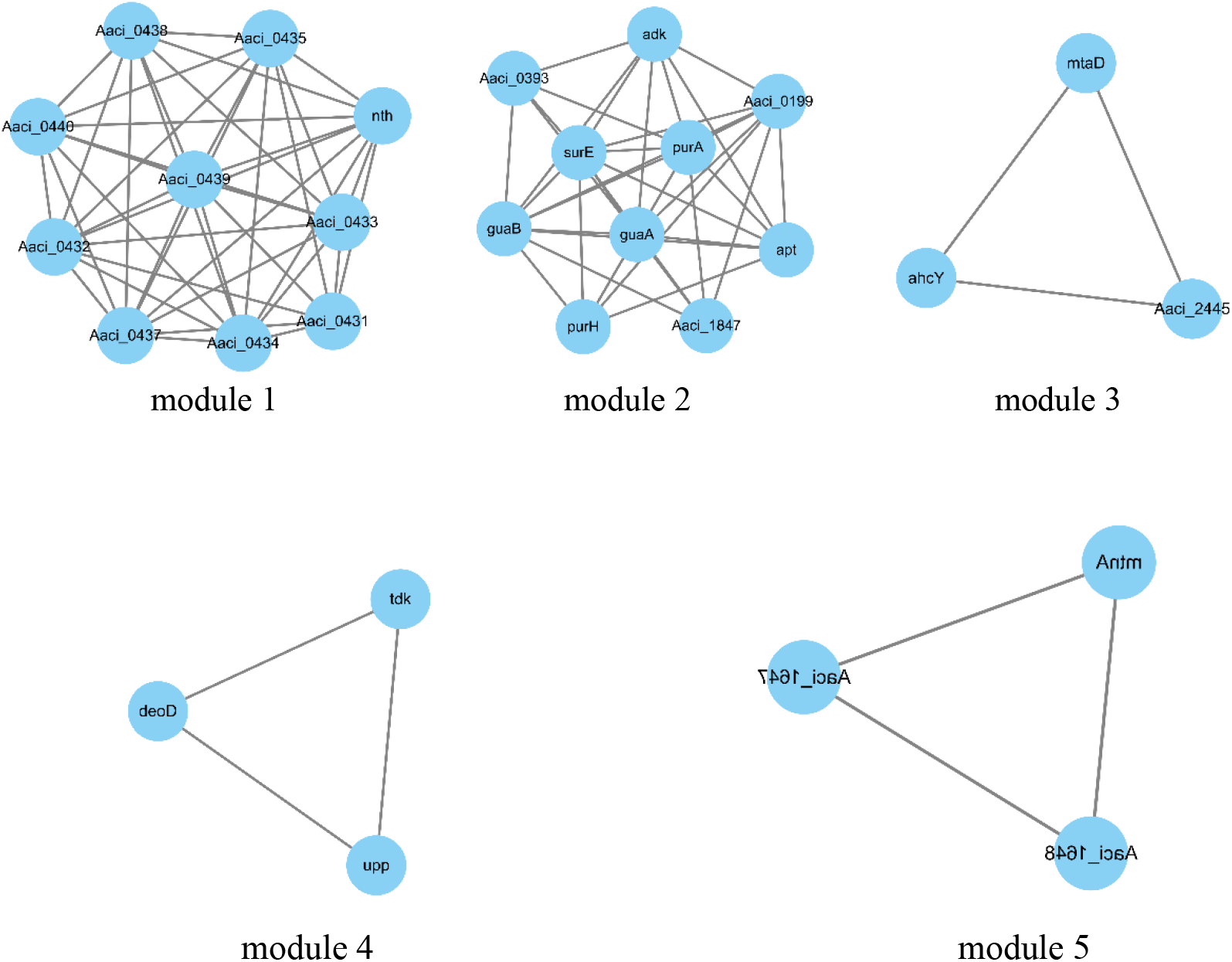
The protein interaction network of inosine.

## Discussion

*A. acidoterrestris* is the major contaminating bacteria in fruit juice production. The acidophilic and heat-tolerant properties of *A. acidoterrestris* and high-stress resistance of its spores pose a serious challenge to the juice sterilization process. Natural bacteriostatic agents not only show bactericidal, bacteriostatic and antioxidant effects but also have the advantage of being non-toxic and harmless. In recent years, the development and application of natural bacteriostatic compounds have become research hotspots in the food processing and preservation field ^[18]^. Inosine is a cellular metabolite of *Bacillus* spp., and also a component of human body. It is widely used in the food and pharmaceutical industries with a variety of medical and healthy applications ^[19]^. It was reported that inosine was a nutrient germination agent of various bacterial spores ^[20-23]^. The effect of inosine on the germination of *A. acidoterrestris* was completely different from the current research (Unpublished data). It is reported that inosine could inhibit the germination of *A. acidoterrestris* spores and cause the death of its vegetative bacteria. Given the unique physiological properties of *A. acidoterrestris*, we speculated that inosine might have a specific bacteriostatic mechanism on *A. acidoterrestris*. Thus, we carried out a series of experiment and find the clues of why inosine could function as bacteriostatic agent against *A. acidoterrestris*.

Previous researches suggest that the bacteriostatic mechanism of natural compounds mainly has three aspects: first, it can destroy cell wall and cell membrane, increase the permeability of the cell membrane which results in the loss of intracellular nutrients and then causes microbial death; second, it can destroy the structure of DNA and RNA which may inhibit gene expression and cause cell death; third, natural compounds can destroy the intracellular mitochondrial structure, resulting in inhibition of microbial respiration and cause hypoxia-induced cell death ^[24, 25]^. In this study, the measurements of surface hydrophobicity, intracellular protein leakage and membrane damage showed that the cell membrane of *A. acidoterrestris* was seriously damaged with the treatment of 5 mM inosines, causing an increase in the permeability of cell membrane and the leakage of nutrients which inhibit the growth and reproduction of the bacterium. However, as a component of the human body, inosine can directly enter the cell through cell membrane, which has the function of promoting hepatocyte recovery and improving hepatocyte activity ^[26]^. The low concentration inosine treatment can cause serious damage to the cell membrane of *A. acidoterrestris*, suggesting that inosine may involve in disrupt cell membrane and exert bacteriostatic effects.

To further study its bacteriostatic mechanism, we applied bioinformatics method to predict the target protein information of inosine and created an interaction network of inosine target proteins. The results showed that the target proteins of inosine were mainly related to DNA cleavage, nucleotide metabolic processes, and purine nucleotide biosynthesis. Genomic DNA is the carrier of genetic information on which cells depend on for their survival and plays a key role in cell growth and development. DNA damage is also a major factor in cell inactivation. DNA lysis activates apoptosis-related signaling pathways that interfere with cell homeostasis and caused apoptosis ^[27, 28]^. Those indicates that inosine might take part in DNA cleavage and damage the cell DNA with its target proteins. Additionally, nucleotide metabolism and purine nucleoside biosynthesis are the core of all biological system metabolic processes. In addition to being closely related to gene replication, transcription, and other biological events, they also show a variety of biological functions in living cells and play an important role in genetic information and energy transmission ^[29, 30]^. In microorganisms, regulatory methods such as attenuation mechanisms, transcriptional repression, feedback inhibition, and feed-forward activation all have an impact on the purine biosynthesis pathway, which in turn affects the supply of energy required for cell reproduction and apoptosis. The target proteins of inosine action are related to nucleotide metabolism, which suggest that inosine might affect cell growth and reproduction via participating in relevant nucleotide metabolic pathway and biosynthesis. It is speculated that inosine might involve in the regulation of the biosynthetic process of purine nucleosides which affects the energy supply required for the growth and reproduction of *A. acidoterrestris*. In addition, the target protein of inosine action may also interact with the translation initiation factor. The translation initiation factor regulates the biosynthesis of proteins which is an indispensable class of proteins in the translation process and plays an important role in various properties of the biological organism. It is found ^[31]^ that down-regulation of the translation initiation factor 4E expression induces apoptosis in breast cancer cells. Therefore, inosine may also inhibit the growth of the bacterium by regulating the activity of the translation initiation factor.

From the analysis above, it can be inferred that the molecular mechanism of inosine inhibition on *A. acidoterrestris* might be caused by regulating nucleotide synthesis and metabolism and translation initiation factor activity. This process can affect DNA replication and cellular energy metabolism and thus inhibit bacterial growth, which needs further investigation and verification. The results of the study analyzed the mechanism of inosine inhibition on *A. acidoterrestris* at the physiological and molecular level, which provides an experimental and theoretical basis for the development of a novel and efficient technique for controlling of *A. acidoterrestris* contamination.

## Acknowledgements

This work was supported by the National Natural Science Foundation of China (Grant number 31771949, awarded to LJ) and the Program for Innovative Research Team in Science and Technology in University of Henan Province (Grant number 21IRTSTHN024, awarded to LJ).

